# Differential Vulnerability of Anterior Cingulate Cortex Cell-Types to Diseases and Drugs

**DOI:** 10.1101/2021.10.26.465972

**Authors:** Marissa A. Smail, Sapuni S. Chandrasena, Xiaolu Zhang, Vineet Reddy, Craig Kelley, James P. Herman, Mohamed Sherif, Robert E. McCullumsmith, Rammohan Shukla

## Abstract

In psychiatric disorders, mismatches between disease-states and therapeutic strategies are highly pronounced, largely because of unanswered questions regarding specific vulnerabilities of different cell-types and therapeutic responses. Which cellular events (housekeeping or salient) are most affected? Which cell-types succumb first to challenges, and which exhibit the strongest response to drugs? Are these events coordinated between cell-types? How does the disease-state and drug affect this coordination? To address these questions, we analyzed single-nucleus-RNAseq (sn-RNAseq) data from the human anterior cingulate cortex—a region involved in many psychiatric disorders. Density index, a metric for quantifying similarities and dissimilarities across functional profiles, was employed to identify common (housekeeping) or salient functional themes across all cell-types. Cell-specific signatures were integrated with existing disease and drug-specific signatures to determine cell-type-specific vulnerabilities, druggabilities, and responsiveness. Clustering of functional profiles revealed cell-types jointly participating in these events. *SST* and *VIP* interneurons were found to be most vulnerable, whereas pyramidal neurons were least vulnerable. Overall, the disease-state is superficial layer-centric, largely influences cell-specific salient themes, strongly impacts disinhibitory neurons, and influences astrocyte interaction with a subset of deep-layer pyramidal neurons. Drug activities, on the other hand, are deep layer-centric and involve activating a distinct subset of deep-layer pyramidal neurons to circumvent the disinhibitory circuit malfunctioning in the disease-state. These findings demonstrate a novel application of sn-RNAseq data to explain drug and disease action at a systems level, suggests a targeted drug development and reevaluate various postmortem-based findings.

## INTRODUCTION

Diseases and clinically relevant drugs have greatly expanded the basic understanding of biology at systems level^1^. A disease and drug are expected to be inversely correlated— loss of homeostasis in disease should be regained by a drug. However, the complexity of cellular networks and variability in cellular vulnerabilities and drug response precludes such simple correlations, making it difficult to understand disease mechanisms and provide effective treatments.

Network complexity and cellular heterogeneity are greater in the central nervous system (CNS) than in other organ systems, and the expected inverse correlation between disease and drug is further reduced in the CNS, as exemplified by a negligible increase in the number of FDA-approved drugs for CNS disorders in the past decade^*1, 2*^. Different CNS cell classes are organized in a dynamic information-processing network, and their differential contribution to the network activity is regulated by many factors, such as their synaptic potentials (excitatory/inhibitory), location (pre-/postsynaptic), and distribution^*3*^. These factors and their variability also contribute to the intrinsic resilience or vulnerability of cells against diseased states. An order of cellular vulnerability has implications in disease mechanisms, as an overly vulnerable cell may represent a critical source of pathophysiology, leading to a domino-like fall of other connected cells in the network^*4*^. Understanding this order will aid in comprehending the complexity of CNS system and serve as a potential avenue for a targeted therapeutic approach across similar diseases.

Finding relative cellular vulnerabilities and drug responses necessitates the comparison of cell-type specific functional profiles across disorders and drugs. For years, this was not possible with whole-tissue samples. Single-cell/nucleus RNA-sequencing (sc/n-RNAseq) now permits analysis of molecular changes at the cellular level. However, most studies employing sc/n-RNAseq are either aimed at characterizing cell-types^*5–7*^ or at answering specific questions about a single disease^*8–10*^. As a result, recent efforts have been limited in their ability to comprehend the canonical relationship between cellular vulnerabilities and drug responses in a disease-state defined by CNS disorders of psychiatric, developmental, and neurodegenerative origin. Where sc/n-RNAseq provides cell-type-specific information, other approaches provide disease- and drug-specific transcriptomic signatures. Useful resources include DisGeNET^*11*^ and the Comparative-Toxicogenomics-Database (CTD)^*12*^, which anchor well curated disease-associated gene-sets, and the connectivity map (cmap)^*13*^, which anchors experimentally derived drug-specific gene-sets. We previously showed that these resources, together with available gene ontologies, can provide insight into shared vulnerabilities across several psychiatric disorders and information regarding druggable-mechanisms for therapeutically manipulating a disease^14^. Here, we hypothesize that systematic integration of human sn-RNAseq and Gene Ontology (GO) with available disease- and drug-specific transcriptomic signatures can 1) differentiate housekeeping and salient events of CNS cell-types, 2) reveal a relative order of cellular vulnerability, 3) identify events undergoing cell-specific changes during disease-states and drug responses, and 4) identify cell-types jointly-participating during disease-states and drug responses Because of their association with several psychiatric disorders^*15–17*^, we analyzed sn-RNAseq data from the anterior cingulate cortex (ACC) of healthy humans to explore these associations. We identified key cellular nodes and events (housekeeping and salient) and mechanism responsible for the observed deviation between drug–disease correlations.

## MATERIALS AND METHODS

An sn-RNAseq dataset corresponding to ACC was downloaded from Allen Brain Atlas (https://portal.brain-map.org/atlases-and-data/rnaseq/human-v1-acc-smart-seq). The data comprise aggregated counts at the gene level collected from 7,283 nuclei using postmortem ACC (left hemisphere) of three adults: 1 female aged (in years) 43 (Postmortem interval (PMI): 18.5 hr; RNA integrity (RIN): 7.4 ± 0.7)) and two males aged 50 years (PMI: 24.5 h; RIN: 7.6 ± 1.0) and 54 years (PMI: 25 h; RIN: 7.7 ± 0.8). The neuronal marker NeuN was used to dissociate and sort the nuclei from individual cortical layers. The collected nuclei were sequenced using SMART-seq v4 RNA-sequencing, and the reads were aligned to exons and introns in the GRCh38.p2 reference genome using STAR aligner. To avoid nuclei clustering based on sex, X and Y chromosomes were excluded from the dataset. The donors had no history of neuropsychiatric or neurological disorders. As such, all cell-specific signatures were regarded as “control.”

Detailed information on data preprocessing, generation of cell-type clusters, filtering cell-specific, vulnerable, druggable, and actionable transcriptomes, theme-centric pathway analysis, density score and disease and drug enrichment analysis is provided in supplementary information.

## RESULTS

### Characterization of ACC-specific cell-types

We investigated an sn-RNAseq dataset from the Allen Brain Atlas (n = 3: male = 2, female = 1) to characterize the cellular composition of ACC. All ACC cells were grouped into 14 distinct clusters (Figure 1A, Supplementary Table 1), which were characterized globally using known cell-type markers (Figure 1B) stratified into cortical layers (Figure 1C), and locally using the gene most enriched in each cluster (Figure 1D).

**Figure 1.**
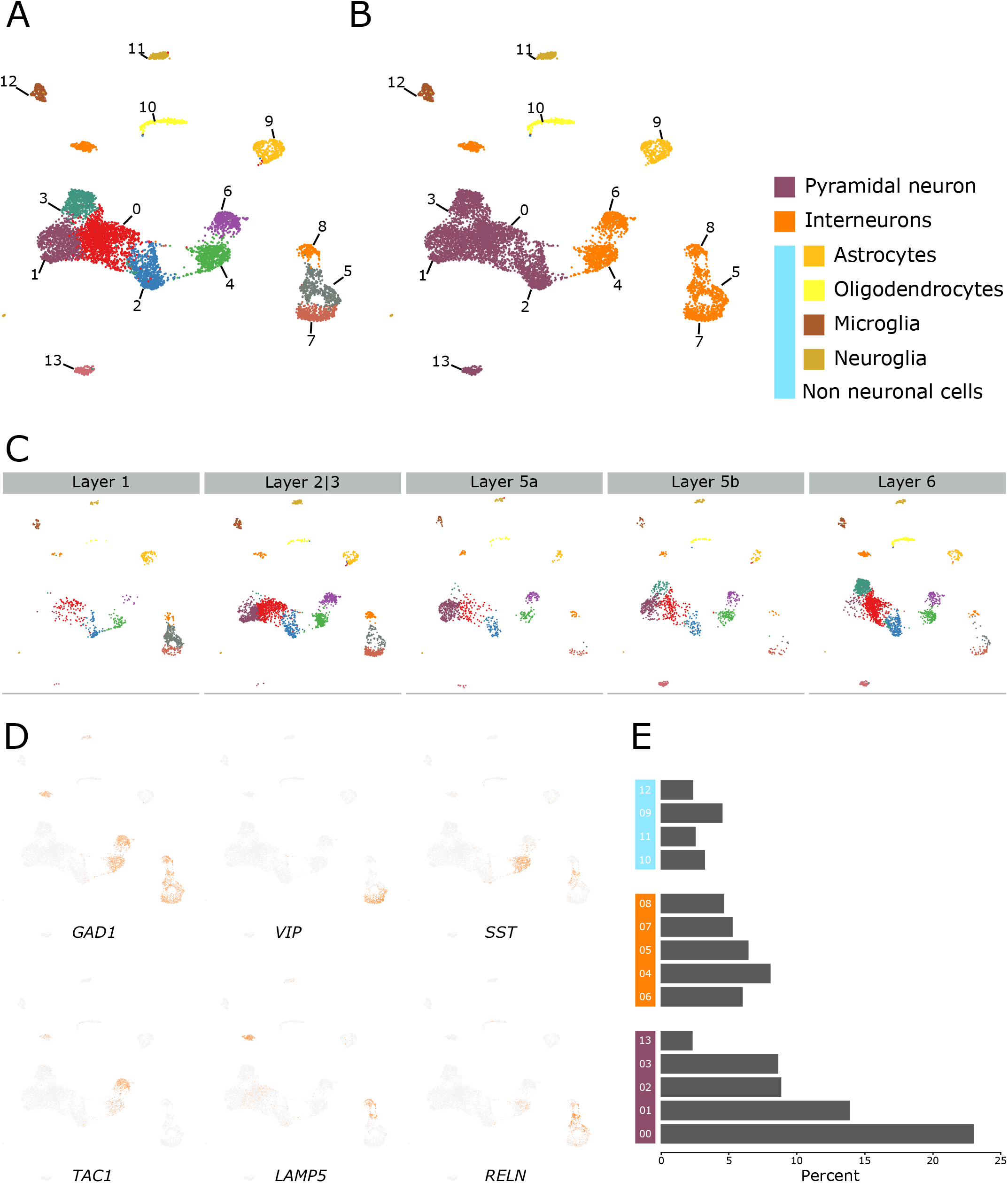
Characterization of control human ACC sn-RNAseq. (**A)** Clustering of human anterior cingulate cortex sn-RNAseq data from Allen Brain Atlas. (**B)** Clusters were characterized using known markers of pyramidal neurons (*SLC17A7*), interneurons (*GAD1* and *GAD2*), astrocytes (*GFAP*), oligodendrocytes (*OPALIN*), microglia (*CX3CR1*) and OPC (*PDGFRA*). (**C)** Cell-type clusters stratified into layers. Notice the enrichment of clusters 05, 07 and 08 in superficial layers 1 and 2|3 and that of cluster 03 and 13 in deep layers 5 and 6. (**D)** Characterization of interneuron clusters using respective markers. See Supplementary Figure 1 for other specific markers. (**E)** Bar plot depicting the proportion of each cell-type cluster.

Interneurons were segregated in the local clustering based on their biological class and origin. *VIP*-, *RELN*-, and *LAMP5*-positive disinhibitory interneurons^*18–20*^ receiving long range^*21, 22*^ and neuromodulatory projections^*23*^ were enriched in clusters 05, 07, and 08, respectively (henceforth: disinhibitory interneuron clusters). These clusters coexpressed corticotrophin-releasing hormone (*CRH*) (Supplementary Figure1) and cluster 05 coexpressed *SST*, *RELN*, and *CRH*. *SST*- and *PVALB*-positive interneurons, originating from the medial ganglionic eminence (MGE)^*24, 25*^, were enriched in clusters 04 and 06, respectively (henceforth: MGE interneurons clusters).

Among the global clusters, the non-neuronal cell-types, including astrocytes (cluster 09), oligodendrocytes (cluster 10), microglia (cluster 12), and oligodendrocyte precursor cells (OPC, cluster 11), exhibited sharper separation without any layer specificity, whereas neuronal cell-types exhibited fluidic boundaries but with layer specificity. For instance, pyramidal neurons (PNs) of clusters 03 and 13 were enriched in deep layers 5b and 6 (henceforth: deep-layer PNs) and disinhibitory interneurons were enriched in superficial layers 1 and 2|3. The remaining PNs (clusters 00, 01, and 02), except those in layer 1, which predominantly anchors interneurons^*26*^, were almost evenly distributed across layers 2|3 through 6 (henceforth: across-layer PNs). MGE interneurons were distributed across layers 1 through 6. Consistent with recent sn-RNAseq data from other areas^*7, 8*^, the three cell classes—PNs, interneurons, and non-neuronal cells— represented 56%, 30.5%, and 12.7% of the total nuclei sequenced, respectively (Figure 1E).

### Functional characterization of ACC cell-types using unique markers

To investigate the functional properties differentiating ACC cell-types, we filtered genes highly (≥2-fold, Supplementary Table 2) and almost uniquely upregulated in each cluster and performed pathway analysis using the GO database. GO terms enriched (*q-value < 0.05*, Supplementary Table 3) across all clusters were categorized into functional themes (Figure 2, left labels). Within different cell-types, across-layer PNs (clusters 00 and 01) were associated with *trans-synaptic signaling*, *channel activity*, and functionalities related to *potassium* ions, whereas deep-layer PNs (clusters 03 and 13), consistent with their long-distance projections^*27–29*^ to other brain areas, were associated with *neuron projection development*, *axons*, and *presynapse*. Interneurons, consistent with their established inhibitory role, were associated with *GABA.* Disinhibitory interneuron clusters, consistent with their long-distance neuromodulatory input^*19, 30*^, were associated with *acetylcholine*, *adrenergic receptor activity*, *catecholamine*, *dopamine*, and *epinephrine*. Non-neuronal cells, consistent with their well-established support and immune-related functionality^*31, 32*^, were associated with *regulation of neurogenesis*, *cell adhesion*, *inflammatory response*, and *innate and adaptive immune response*. *Cell death* and *homeostatic process* were associated almost exclusively with non-neuronal cells.

Density scores (see methods; Figure 2, right), an index to quantify how common (close to 1) or unique (close to 0) a theme is across different cell-types, were generated for each theme. Common and unique themes also reflect housekeeping and salient functionalities, respectively. Themes associated with neuronal structure (*neuron projection development*, *axon*, *dendrites*, *trans-synaptic signaling*, *synapse organization*, and *presynapse*), cell signaling (*intracellular signal transduction* and *regulation of protein phosphorylation*), *vesicles*, and *extracellular region* had high densities (>75th percentile, Figure 2, in red), suggesting that these functions are the collective result of coordinated efforts among multiple cell-types. Themes associated with *proteolysis*, immune response (*inflammatory response*, *adaptive immune response*), neurotransmission (*acetylcholine*, *adrenergic*, *dopamine*, *epinephrine*, and *serotonin)*, and ions (*calcium* and *sodium*) had low densities (<25 percentile, Figure 2), suggesting that these functions are more distinctive to specific cell-types. All low-density salient functionalities were associated with either disinhibitory interneurons or non-neuronal cells.

Overall, unique cell-type-specific markers in a “healthy” (control) state can precisely identify unique functional properties associated with ACC cell-types and can segregate housekeeping and salient functionalities associated with them; thus, giving confidence to integrate and build new ontologies around them.

### *VIP* and *SST* interneurons: most vulnerable cell-types

To infer relative cellular vulnerabilities, we reasoned that a vulnerable cell would tend to have the highest association with CNS disorders of different origins (complex, mendelian, or environmental). We searched for statistical enrichment of experimentally curated CNS disorder-specific gene-sets from DisGeNET in our cell-type-specific markers. In total, 112 CNS-disorder-associated gene-sets, categorized into psychiatric-, developmental-, neurodegenerative-, and other CNS disorders, were enriched (*p-value* < 0.05; Figure 3A; Supplementary Table 4) in at least one cell type. Interneurons were implicated in the highest number of diseases (orange; ~30%), followed by non-neuronal cells (blue; ~15%), and PNs (purple; ~10%). *VIP* and *SST* interneurons (clusters 05 and 04, respectively) were the most implicated, whereas across-layer PNs (cluster 02) were the least implicated. Among non-neuronal cells, astrocytes (cluster 09) and microglia (cluster 12) were implicated in a relatively high number of diseases.

**Figure 3.**
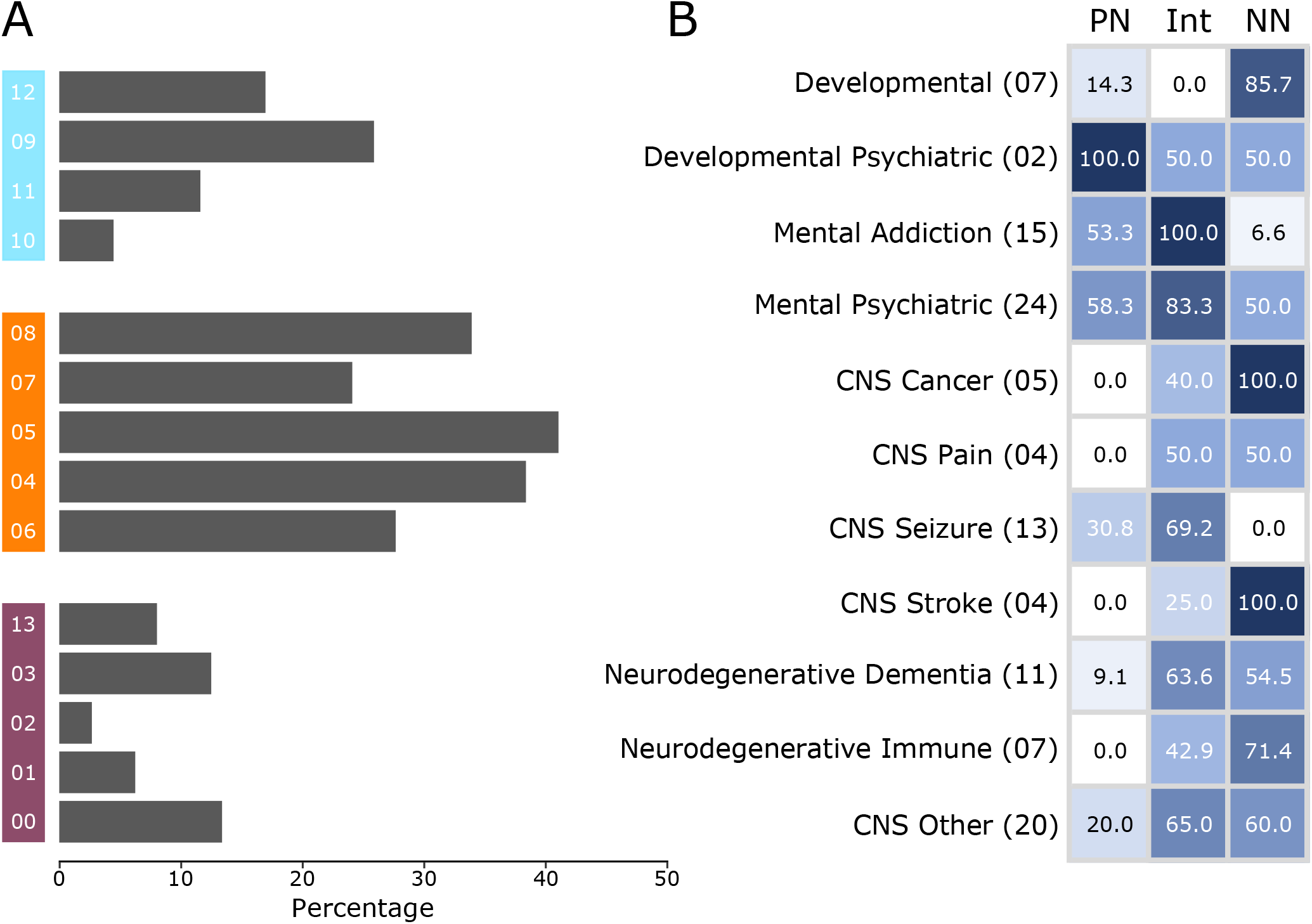
Distribution of CNS disorders by cell-specific enrichment. **A)** The percentage of CNS disorders (out of 112) that have significant (*p-value* <0.05) gene-set overlap with a specific cell-type cluster. Cluster numbers are provided at the left, with non-neuronal clusters in blue (top), interneuron clusters in orange (middle), and PN cluster in purple (bottom). **B)** Percentage, of disease categories associated with PNs, interneurons (Int), and non-neurons (NN). Abridged disease categories (left labels) as provided by DisGeNET are used. The lighter-to-darker shades of blue represent the percentage of disease categories associated with each cell class. The number of diseases in each category are provided in parentheses. See Supplementary Figure 2 and Supplementary Table 4 for more details.

In terms of disease types (Figure 3B; Supplementary Figure 2, Supplementary Table 4), interneurons were implicated in nearly all CNS disorders and comorbid conditions, including seizures and dementia. There was no clear segregation of disorders based on MGE and disinhibitory interneurons, but some disorders related to addiction (*amphetamine-related disorder*), psychiatry (*schizoaffective disorder*), *neurogenic inflammation*, and *generalized hyperkinesia* were preferably associated with MGE interneurons. Certain disorders related to psychiatry (involving cognition and behavior), cancer (*malignant neoplasm*), seizure (*West syndrome*), and neurodegenerative disorder (*Parkinson’s disease*) were preferably associated with the disinhibitory neurons. Non-neuronal cells, consistent with their proliferating nature^*33*^ and immunological functions^*31, 32*^, were most enriched in developmental, cancer, and neurodegenerative disorders. PNs had fewer overall associations, but those present across all layers (clusters 00, 01, and 02) were strongly implicated in seizure (*epilepsy*, *petit mal*, and *status epilepticus*) and addiction (*Alcohol intoxication*, *cocaine abuse*, and *substance withdrawal syndrome*), whereas those dominant in deep layers (clusters 03 and 13) were implicated in developmental disorders (*language delay and autistic disorders*) and drug-induced psychosis.

Overall, the overlap of cell-specific markers with disease signatures reveals that cell-types exhibit some specificity for different disease groups. Among different cell classes, interneurons, particularly *SST-* and *VIP-*positive have the highest vulnerability to disease, followed by non-neurons and PNs.

### Vulnerable transcriptome revealed cell-type-specific functional changes in disease-states

Present data collected from healthy individuals prevents the study of cell-specific functional responses in disease-states. To circumvent this, we pooled the genes that drive the enrichment (i.e., significant overlap) of 112 disease-specific gene-sets defined above in each cell-type-specific gene-set (i.e., leading-edge genes^*34*^; Supplementary Figure 3, blue). Cell-type-specific pooled gene-sets constitute the “vulnerable transcriptome,” (Supplementary Table 5) and the 112 disorders represent the “disease-states.” To identify altered functionality of a cell type in the disease-state, the pathway profiles associated with each cell-type-specific vulnerable transcriptome (*q-value* < 0.05, Figure 4, blue, Supplementary Table 6) were observed against the backdrop of the control cell-specific pathways (Figure 4, yellow). A positive shift in density score (Figure 4, red vectors, see legend), implying a theme-centric increase in associated cell-types or pathways, indicates themes vulnerable in disease-states.

**Figure 4.**
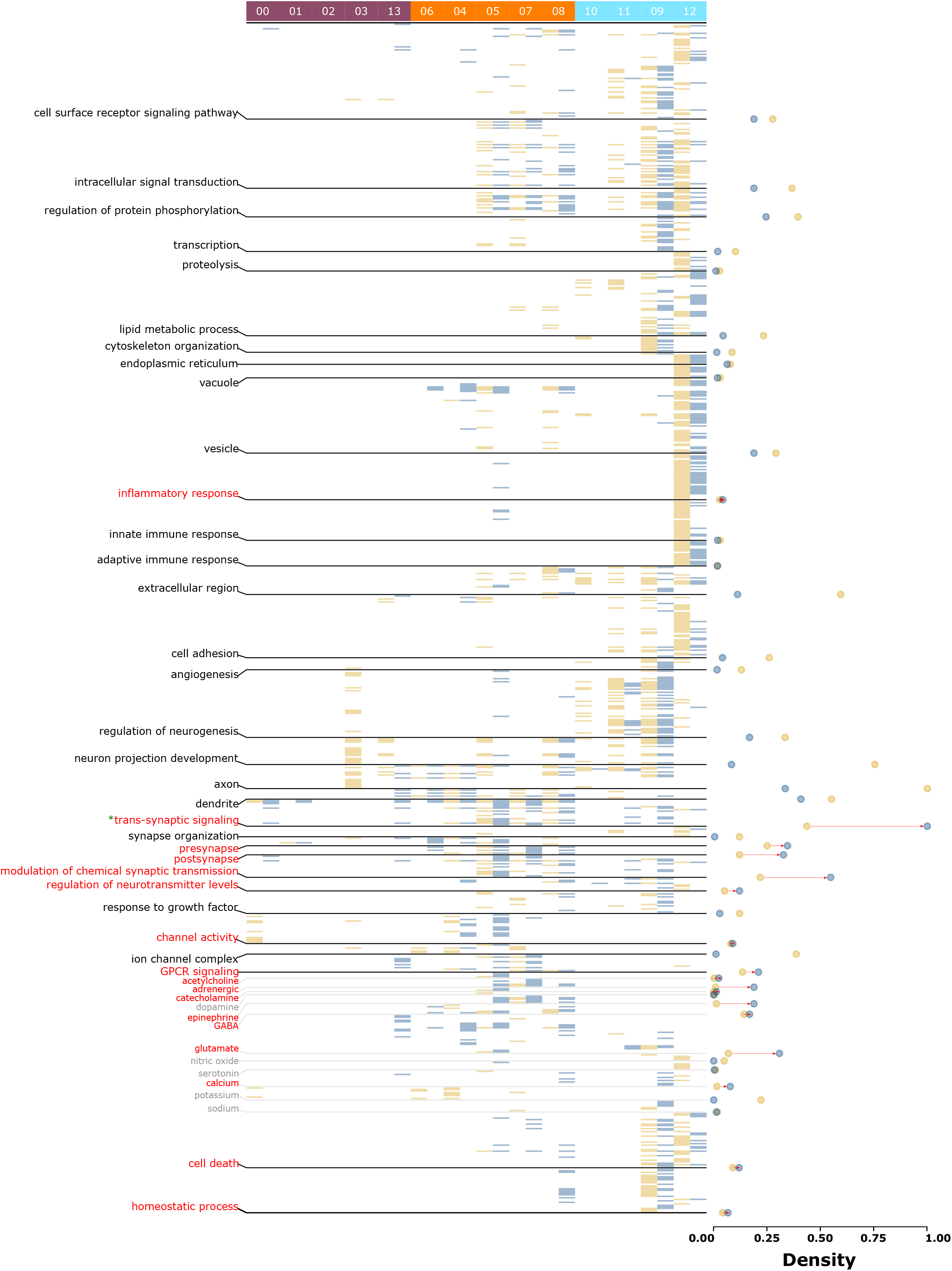
Cell-type specific functional state comparison between control and disease-states. Heatmap of significant (*q-value* <0.05) pathways for each cell-type specific gene-set and vulnerable transcriptome representing control (yellow) and disease-state (blue), respectively. Cell-type cluster numbers are provided at the top, with PN clusters in purple (left), interneuron clusters in orange (middle), and non-neuronal clusters in blue (right). Pathways are organized by functional themes provided at the left. Each state’s theme-densities are represented by solid circles of the respective colors at the right. Note that an increase in disease-state density (marked with red vectors) represent vulnerable themes involving additional cell-type or pathways. However, note that, as the directionality (up- or downregulation) of the disease-specific gene-sets is unknown, an increase in density only reflects the theme’s involvement in the disease-state and not its upregulation. As the cell-specific densities (yellow) cancel out the low densities in the vulnerable state (blue), the decreased density has no greater (or lesser) influence on the cell’s specific theme than the cell’s baseline state. See Supplementary Table 12 & 13 for more details.

Inflammatory, synaptic (*trans-synaptic signaling*, *presynapse*, *postsynapse*), neuromodulation (*calcium*, *acetylcholine*, *adrenergic*, *catecholamine*, *epinephrine*, *GABA*, and *glutamate*), neuromodulatory infrastructure (*modulation of chemical synaptic transmission*, *regulation of neurotransmitter levels*, *channel activity*, *G protein-coupled receptor signaling pathway*), cell death, and homeostatic changes were vulnerable themes across the cell-types. Except for the housekeeping themes involving *trans-synaptic signaling* (Figure 4, “*”), the disease-state largely influenced low-to-mid-density themes, salient in fewer cells or cell classes in control conditions, yet increasingly common in disease conditions. Within cell-types, across-layer PNs (clusters 00 and 01) showed increases in trans-synaptic signaling-associated pathways. Among deep-layer PNs (clusters 03 and 13), the two clusters showed an opposite effect. Cluster 03, highly enriched in the control-state, was not affected in the disease-state, whereas cluster 13, not influenced in the control-state, was associated with new themes: *dendrites*, *trans-synaptic signaling*, *presynapse*, *modulation of chemical synaptic transmission*, *GPCR signaling*, *glutamate*, and *calcium* ion-related events. Among interneurons, those from MGE (particularly *SST*-positive cluster 04) in the disease-state were enriched in *cell-surface receptor signaling pathways*, *vesicles*, *cell adhesions*, *regulation of neurotransmission level* and *glutamate*-related activity; those related to disinhibitory interneurons in the disease-state showed additional enrichment in *inflammatory response*, *innate immune* response, *regulation of neurogenesis*, *cell death*, and *homeostatic process*. Notably, these themes in the control state exclusively belonged to non-neuronal cells. Among non-neuronal cells, in disease-state, astrocytes (clusters 09), exhibited increase in pathways associated with *trans-synaptic signaling, modulation of chemical synaptic transmission*, *regulation of neurotransmitter levels*, and signaling associated with *glutamate* and *calcium.*

Overall, a direct comparison of control and disease-states reveals that disease-state transitions are biased toward cell-specific salient themes and entail the addition of new biological functionalities or an increased involvement of more cell-types. The new functionalities were biased toward the neuron localization and functionality. The cell-types distributed across layers (PNs, MGE interneurons, and non-neuronal cells) were associated with all aspects of signaling (intracellular and synaptic) and neuromodulatory events, whereas the disinhibitory cell-types enriched in superficial layers were associated with immunological and homeostatic functions.

### Deep layer pyramidal neurons are most responsive to drugs in disease-state

Next, we reasoned that, like enrichment of disease signatures in cell-type specific gene-sets, enrichment of drug signatures in cell-type specific gene-sets would identify druggable transcriptomes (Supplementary Figure 3, maroon; Supplementary Table 7) indicating drug impact in the control state (henceforth: druggable state). Likewise, enrichment of drug signatures in cell-specific vulnerable transcriptomes identified above would reveal actionable transcriptomes (Supplementary Figure 3, green; Supplementary Table 8) indicating drug impact in the disease-state (henceforth: actionable state). For both states, a cell-specific drug mechanism of action (MOA, Supplementary Figure 4; Supplementary Table 9) and pathway profile similar to that described in the disease-state can be investigated.

In the druggable state drugs have a greater impact on non-neurons and interneurons, but not on PNs (Supplementary Figure 4A, maroon). Surprisingly, however, in the actionable state (Supplementary Figure 4A, green), fewer drugs impacted astrocytes and microglia, but more drugs impacted the disinhibitory interneurons and deep layer PNs. The drug MOA (and target, Supplementary Figure 4B) associated with actionable state shows that among the three classes, PNs are targeted through neurotransmitter system (*acetylcholine-, dopamine- and histamine-receptors and serotonin-reuptake*), hormones (*androgen- and estrogen-receptors*) and DNA metabolism (*DNA-replication and synthesis*); interneurons, are targeted through kinase system (*ABL, BCR-ABL, and FLT3 kinases*), neurotransmitter-system (adrenergic- and serotonin-receptors) and growth factors (*EGFR*) and non-neuronal cells are targeted through enzymes (*topoisomerase, HMGCR, HDAC, cyclooxygenase ATPase*) and structural components involving *tubulin*, and *angiotensin.*

With respect to pathway profiles (*q-value* <0.05; Supplementary Table 10&11), similar to the disease-state, the druggable-(Supplementary Figure 5, red) and actionable-(Supplementary Figure 6, green) states displayed an increase in density associated with inflammatory response, neuromodulator activity (*adrenergic, catecholamine, epinephrine*), events involving calcium ions, cell death and homeostatic process. However, unlike the former the latter two displayed an increased density associated with themes involving signal transduction (*intracellular signal transduction and protein phosphorylation regulation*), organelles (*endoplasmic reticulum and vacuole*), adaptive immune response, serotonin-related neuromodulation, and sodium ion related activity. Interestingly, the actionable transcriptome influenced greatest number of housekeeping themes (4/11, Supplementary Figure 6, green “*”) when compared to both the disease (1/11) and druggable state (2/11).

To understand the functionality associated with the actionable state, we directly compared its pathway profile with druggable state (Supplementary Figure 7 notes). Here, a positive shift in theme-density (Supplementary Figure7, red vectors) implies increment in cell-types or pathways in association with actionable state. Reflecting an increase in activity of associated neurons, the increased theme densities of actionable state involved components of Excitation/Inhibition balance (i.e., Glutamate and GABA) and action potential (calcium, potassium, sodium, and ion channel activity).

Overall, the segregation between cell-specific druggable- and actionable-state shows that drugs associations are stronger when disease is considered and PNs, which are barely implicated under control and disease-state, have the most substantial impact, largely by means of activating the deep layer PNs. Furthermore, actionable state, unlike the disease-state, preferably targets the housekeeping themes.

### Functional profile clustering captures nonrandom associations between ACC cell-types

GO terms represent coordinated expression^*35, 36*^ of genes across different biological-processes, molecular-functions, and cellular-components. In this regard, a GO term’s enrichment in two or more cell-types originating from the same tissue but defined by unique gene-sets indicates a joint-participation of these cell-types against the enriched GO terms (Supplementary Figure 8 notes). Accordingly, the functional similarities between the cells governed by the cell-type-specific markers of a given state indicate their joint-participation against some afferent input or efferent output^*37, 38*^ associated with that state. Based on this proposition to better understand the joint participation of cell-types in the control, vulnerable, druggable, and actionable states, we clustered the cell-types based on their entire functional pathway profiles (*q-value* <0.05; Supplementary Table 12&13) associated with the respective state and searched for studies that experimentally validated the results for control and disease-state (Figure 5). In the control state, the PNs, consistent with our previous report^*4*^ and concurrent receipt of long-range input along with the *VIP* interneurons, clustered closely with disinhibitory interneurons^*18, 39*^. Among the non-neuronal cell-types, consistent with their coupling through gap junctions^*40*^ and joint participation to deliver energy substrates for sustained neuronal firing^*41*^, astrocytes, oligodendrocytes, and OPCs were grouped together in all the states. Consistent with the increased activity of astrocytes in pathophysiological states^*42–45*^, an increased functionality of astrocytes (see circle size) in states other than the control state is apparent. Increased microglia interaction with PNs in the prefrontal cortex in psychiatric diseases^*46, 47*^ and other CNS-disorders^*48*^ has been reported. Consistent with these findings, we observed a subset of deep-layer PNs (cluster 03) jointly participating with microglia in the disease-state.

Overall, functional clustering of cell-types in the control and disease-states points to the functional activity and nonrandom joint participation of cell-types in different physiological brain states, most likely reflecting the altered routing of information (afferent input and efferent output) in these states.

## DISCUSSION

The specific impact of diseases and drugs on different CNS cell-types remain poorly understood. Using cell-type-, disease-, and drug-specific signatures, we segregated the cell-specific vulnerable, druggable, and actionable transcriptomes of ACC cell-types and developed ontologies around the disease, druggable, and actionable states, respectively. Our approach has several advantages over sn-RNAseq in a case–control setting. First, a signature-based approach is used to detect changes in the disease-state. These signatures, curated from various resources, summarize the collective biological state of a phenotype^*49*^ and enable direct comparison across several other phenotypes, thereby minimizing various technical challenges associated with data normalization and regressing disease-specific confounding and latent effects^14^. Second, the disease-state is defined using psychiatric, developmental, and neurodegenerative CNS disorders that overlap significantly with the cell-specific signatures. This enabled investigation of the vulnerable canonical microcircuitry elements and modules across the disease-states and, for the first time, objectively identify a hierarchy of cellular vulnerability. Third, the approach, also for the first time, contrasted the drug and disease response and as discussed in the subsequent sections, revealed that they act on different cell-types, anatomical layers, microcircuitry modules, and downstream effector areas. Fourth, functional pathways, specifically from the GO database, are highly redundant. This redundancy reflects biological parsimony, in which a single gene may participate in multiple similar biological processes. We used the density index, which compresses redundancy and leverages biological parsimony, to compare the cell-specific functional profile of any two states. Finally, by leveraging the notion that shared pathways between cell-types demonstrate coordination, we identified cell-type clusters (Figure 5) jointly participating in processing the altering information in each state. Interesting patterns of cellular response and interaction emerged, many of which are supported by previous experimental studies, lending credence to the validity of our approach.

**Figure 5.**
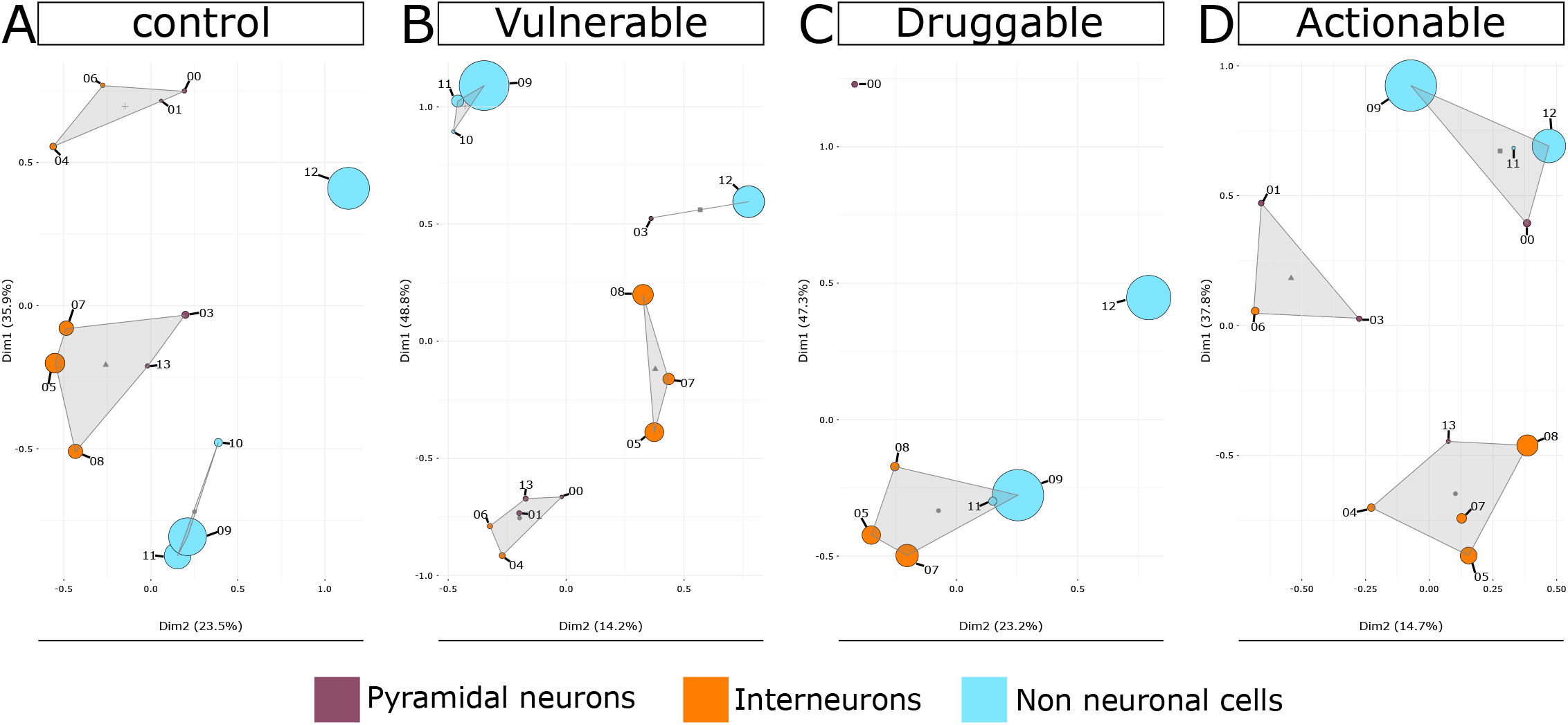
Functional clustering of cell-types in control, disease, druggable, and actionable state. Principal component-based clustering of the complete GO terms profiles associated with each cell-type cluster (shown as circles) in **A)** cell-type specific control-state, **B)** disease-state, **C)** druggable-state, and **D)** actionable-state. Cell-type clusters are labeled by their respective numbers as shown in Figure 1. Non-neuronal clusters are depicted in blue, interneuron clusters are depicted in orange, and PN cluster are depicted in purple. The size of each circle is proportional to the number of pathways associated with the cell-type cluster. Circles that are closer in space exhibit stronger functional similarity, with triangles indicating the centroid of the functional cluster.

### Disease-states are superficial layer-centric and involve alterations in disinhibitory neurons

In the control state, as defined by cell-specific signatures, we identified themes that are common (housekeeping) or unique (salient) between ACC cell-types. Comparison between the control and disease-states (Figure 4) revealed several interesting changes at the cellular level. First, disease-state transitions are biased toward cell-specific salient themes and entail the addition of new biological functionalities or an increased involvement of more cell-types, suggesting a functional reorganization of cellular processes. Second, interneurons were most vulnerable (Figure 3), and, among interneurons, *SST* and *VIP* neurons were associated with most diseases. Their functionality in cortical microcircuits suggests that dendritic inhibition [a function of *SST* interneurons^*50, 51*^], and disinhibition [a function of *VIP* interneurons^*52–54*^] are common circuitry modules that are altered during the disease-state. Third, the disinhibitory neuron subgroups distributed in superficial layers exhibited additional functionalities largely associated with homeostatic functions (Figure 4), which, in the control state, are linked to astrocytes and, to some degree, to microglia. This could be associated with some receptors common between disinhibitory interneurons and astrocytes but are expressed in the interneurons upon an aversive stimulus. This notion, as detailed next, is further supported by the joint participation of astrocytes and disinhibitory neurons in the druggable state, where astrocytes are most influenced (Figure 5). The disease-specific homeostatic gain of the disinhibitory neurons is also supported by their joint participation in the disease-state, perhaps linked with their involvement with *CRH* involved in homeostatic balance^*55*^ and exclusively co-expressed in these neurons (Supplementary Figure 1). Fourth, shifting the established coordination between the superficial layers’ disinhibitory neurons (Figure 5A) and two groups of deep-layer PNs known to have dendritic projections to superficial layers, we observed joint participation between microglia and a subset of deep-layer PNs in the disease-state (Figure 5B). The other subset of PNs as discussed next, was involved in drug action. Overall, mapping these vulnerable functionalities to the layer-specific distribution of the neurons suggests that most of the vulnerable action occurs in the superficial layer, where the information from different areas is merged^*56, 57*^. These actions are largely influenced by homeostatic processes and exhibit a shifting balance of input coming to deep-layer PNs and disinhibitory interneurons.

### Drug actions are deep layer-centric and involve alterations in PNs

We explored the drug action at two levels: the druggable state, representing drug effects without disease, and actionable state, representing drug effects with disease. In the druggable state, drug effects appear to mimic the disease-state by representing a drug’s residual effect following a disease or a drug’s side effect. A homeostatic alteration can be expected in both cases^*58*^. Supporting this, we observed that similar to disease-state, the drug action only targets the salient themes and has the highest involvement of astrocytes and microglia (Supplementary Figure 4), which are predominantly linked to the homeostatic process. In contrast to the disease-state, however, we observed joint participation between disinhibitory interneurons and astrocytes (Figure 5C). This suggests that unlike in diseases, drugs may alter the aforementioned receptors shared by both cell-types, most likely through volume transmission-based mechanisms recently reported to activate shared recurrent synaptic circuitry in these cell-types^59^. In a similar vein, we recently studied the effect of remission from major depressive disorder in the presence of drug effect^*60*^. Corroborating with this finding, we found that the superficial layers of disinhibitory interneurons showed the highest association with non-neuronal cells.

In the actionable state, encouragingly, the drugs show much stronger effects and following observation were of note. First, unlike in the disease- and druggable states, housekeeping pathways distributed across several cell-types were most affected (Supplementary Figure 6). This agrees with the notion of drug promiscuity in neurological disorders where the drugs simultaneously influence many cell-types^*61*^. As our results indicate, the influence primarily entails transient cellular activities and an altered E/I balance. The exaggerated involvement of PNs (Supplementary Figure 4), as opposed to the more vulnerable interneurons or non-neurons, suggests that the shift in the E/I balance is more toward excitation. Second, deep-layer PNs (cluster 03) jointly participating with microglia in the disease-state differ from the deep-layer PNs (cluster 13) exhibiting an exaggerated activity in the actionable state. Considering the division of deep-layer PNs into intratelencephalically and extratelencephalically projecting subtypes^*62*^, the nonoverlapping targets of diseases and drugs demonstrate a disparate downstream effect. Third, considering the MOA associated with the overrepresented PNs in the actionable state (Supplementary Figure 4), we observed an overly represented monoamine-related effect. In contrast to the specific association of disinhibitory neurons with the monoaminergic afferents in the control state, the overrepresentation of monoamine-related effects in PNs suggests that the abnormal disinhibitory effect in the disease-state is bypassed by using drugs. Overall, our results suggest that drugs in action, rather than blocking the initial cause of the disease, serve as a bandage on downstream processes, likely ameliorating the disease symptoms rather than resolving the initial problem.

### Endophenotypes of drug action

The observed joint participation of disinhibitory interneurons and non-neuronal cells in the druggable state (Figure 5C) and the prominence of layer 5 PNs in the actionable state (Supplementary Figure 4) reflects several postmortem studies showing the effect of interneurons and astrocytes^*63, 64*^ as well as layer 5 specific^*65–67*^ effects in psychiatric disorders. While these studies posit that these effects are linked to disease effects, our results suggest that they may be either an artifact or an important endophenotype of drug action. While a few studies support this conclusion^68, 69^, we established it independently in schizophrenia.

### Limitations

Our approach has some inherent limitations. First, we aimed to identify inconsistencies between the disease-state and drug response at systems level, with disease and drug response defined using a range of CNS disorders and drugs, respectively. Experimental validation is difficult unless individual disorders are evaluated, which would contradict the purpose of identifying conserved microcircuitry motifs and vulnerable cell-types. However, these findings can be incorporated into a biophysically realistic computer model^*70*^. Second, the data were acquired from the nucleus, which limits the number of pathways studied due to its poor coverage. This includes information on mitochondrial function, which can be affected by drugs and diseases. Third, the analyses were based on upregulated genes for each cluster, preventing us from drawing conclusions on processes downregulated in disease or drug states.

## Acknowledgments

**Funding**: MAS is supported by National Institute of Mental Health predoctoral fellowship F31MH125541.

**Author contributions**: R.S. and M.A.S. conceptualized the study and together wrote the manuscript. S.S.C. participated in formulating the density score. X.Z., V.R., and R.E.M., participated with R.S. and M.A.S. in developing drug-based ontology. C.K., M.S., and J.P.H. participated with R.S. and M.A.S. in developing disease-based ontology. All authors participated in writing the manuscript.

**Data and materials availability**: All data are available in the main text or the supplementary materials.

**Conflict of interest**: The authors declare no competing interests. Supplementary information is available at MP’s website.

## Figure legends

**Figure 2. Functional analysis of cell-types in the control state.** Heatmap of significant (*q-value* <0.05) pathways for each cell-type. Cell-type cluster numbers are provided at the top, with PN clusters in purple (left), interneuron clusters in orange (middle), and non-neuronal clusters in blue (right). Pathways are organized by functional themes provided at the left. Densities for each theme are provided at the right, with higher indices (in red, > 75^th^ percentile) indicating more common (housekeeping) processes and lower indices (in yellow, < 25^th^ percentile) indicating more unique (salient) processes. The vertical line at density = 0.26 represents the average density across all themes. See Supplementary Table 3 for more details.

