## Supplementary Information for "Differential Vulnerability of Anterior Cingulate Cortex Cell-Types to Diseases and Drugs"

### MATERIALS AND METHODS

#### Data preprocessing

The count matrix and barcode data were loaded into the R programming language through the *Read10X* function in the Seurat R package. A usual quality control step in the analysis involves, removing low-quality cells having very few genes (<200), cell doublets identified by aberrantly high gene counts (>2500), and dead cells identified by excessive (>5%) expression of mitochondrial genes. All the 7,283 nuclei passed these criteria. The counts were normalized to 10,000 counts per cell and transformed to log scale using the default global-scaling normalization implemented using *NormalizeData* function in Seurat. To ensure that the cell-specific marker genes identified were not influenced by PMI—a factor greatly influencing postmortem studies[1], we included them in the metadata and then regressed them with *ScaleData* function. The average RIN (~7.6) of all samples met the quality requirements for gene expression research[2, 3].

#### Generation of cell-type clusters

To highlight biologically relevant cell-type clustering, genes exhibiting high cell-to-cell variation in the datasets [i.e., highly variable genes (HVGs)] play an important role[4]. With default selections and cutoffs, the *FindVariableGenes* function in Seurat was used to identify 2000 HVGs, accounting for the top 10% of the scaled dispersion in the data. Principal component analysis was performed on HVGs and based on the elbow plot, which ranks the PCs based on percentage variability explained. The first 20 PCs were used to generate cell-type clusters using the *FindClusters* function. The function includes a resolution parameter (typically ranging from 0.4 to 1.2) that specifies the degree of granularity of the cell-type clustering, with increasing values resulting in a greater number of clusters. Unfortunately, the clustering algorithm generates an arbitrary number of clusters, and there is no consensus regarding how to adjust the resolution[5]. An increasing number of clusters (i.e., higher resolution), however, splits the existing cell-type cluster into a possibly new subtype, whereas a decreasing number of clusters (i.e., lower resolution) can better characterize the existing cell-types. We looked at the broadest layer-specific clustering of known cell-types; accordingly, a resolution of 0.6 was selected, which generated 14 clusters visualized using a uniform manifold approximation and projection (umap) plot (Figure 1, Supplementary Figure 1a &1b). The clusters were characterized based on the expression of canonical markers of inhibitory neurons (*GAD1*, *GAD2*, *SST*, *PVALB*, and *VIP*), PNs (*SLC17A7*), astrocytes (*GFAP*), microglia (*CX3CR1*), oligodendrocytes (*OPALIN*), and OPC (*PDGFRA*). Cluster-specific enriched genes i.e., all genes differentially expressed and upregulated in the cluster of interest with respect to the remaining clusters, were identified using the *FindAllMarkers* function with default settings (Supplementary Table 1). The data were already stratified into layers. Therefore, post-hoc layer identification or layer-specific cell-type enrichment was not performed. However, to observe the distribution of cells around different layers (Figure 1C), the *split.by* function was used.

**Filtering cell-specific, vulnerable, druggable, and actionable transcriptomes**

Cell-type enriched markers identified by Seurat can only distinguish cellular functions at a broad level shared by many cell-types in the same class (PNs, interneurons, or non-neuronal cells). Conversely, cell-type-specific markers engineer cell functional properties[6] and are essential to understand the unique properties of cellular subtypes within a cell class. Thus, to identify cell-type markers with greater specificity, we filtered the Seurat output so that a gene expressed in a cell-type cluster of interest has a minimum average log fold change expression of ≥0.25 and is at least twice more than the remaining clusters. Applying more stringent rules, such as increasing the average minimum fold change difference, excluding genes coexpressed with other cell-types, or increasing the fold change difference between the cluster of interest and the remaining clusters, significantly reduces the number of marker genes and their related functional profiles.

Key input to any functional analysis is unbiased gene-list summarizing a biology phenomenon or a phenotype[7]. Based on this premise, we reasoned that cell-type-specific genes-sets statistically (see enrichment analysis below) overlapping (Supplementary Figure 3) with disease- and drug-specific gene-sets can reveal the vulnerable and druggable subsets (of cell-type specific genes-sets), which can not only explain the functional contribution of each cell type to the disease and druggable states, respectively; but can also enumerate diseases a cell type responds to (Supplementary Figure 2) and the MOA associated with each cell type (Supplementary Figure 4). Furthermore, the overlap between the vulnerable and drug gene-set can reveal subsets of genes that respond to drugs in disease-states (referred to as actionable) and reveal functional contributions and drug MOA in response to the disease-state. The disease-specific expert-curated gene-sets were downloaded from DisGeNET (https://www.disgenet.org/downloads)[8], and the drug-specific gene-sets were downloaded from drug signature database (DSigDB[9]); http://dsigdb.tanlab.org/DSigDBv1.0/. DSigDB contains gene-sets from various sources, including that from cmap[10], and CTD[11]. Due to the study's objective of elucidating the MOA linked with a particular cell type in a disease-state, only drugs with available MOA were considered. MOA for drugs were downloaded from https://clue.io/touchstone and http://ctdbase.org/downloads/

**Theme-centric pathway analysis**

All the gene-sets from different states (control, vulnerable, druggable, and actionable) were employed to perform pathway analysis using GO. All pathway profiles ($q\text{-}value < 0.05$) were tagged for their respective cell type and state and stacked vertically in to one file. The enrichment score (ES) for each pathway was determine by $-\log_{10}$ transformation of the q-value (i.e., $-\log_{10}(q\text{-}value)$). To compare ES across different cell-types and states, the vertically stacked pathway profile was pivoted using the *group.by* function in R (Supplementary Table 12). To summarize and filter the large number of pathways across all cell-types × state combinations to neurobiologically relevant themes, we leveraged the hierarchical organization of the GO database and picked 44 GO terms (themes) belonging to biological process-, molecular function-, or cellular component-related ontologies representing most neurobiological functions of interest and then looked for the GO terms in our results, which are linked to these 44 themes as child nodes using *GOBPANCESTOR*, *GOMFANCESTOR*, and *GOCCANCESTOR* functions in R *GO.db* package in R. The theme summarized pathways across cell-type × state combination (Supplementary Table 13). The ES of all pathways across all cell-types × state combinations were clustered (Figure 5) using the *FactoMinerR* package in R.

**Density score**

To quantitatively differentiate how common or unique a theme was across different cell-types × state combinations, we modified the density score introduced in our earlier work[12]. A theme $A_{ts}$ represents a matrix embedded within a larger *t* × *s* matrix, where $t = 1,2,\ldots.,44$ represents the themes and $s = 1,2,3,4$ represents the states. Within the matrix $A_{ts}$, an element can be represented as:

$$A_{ts} = (a_{pc})_{p=1,2,\ldots.r_t, c=1,2,\ldots.14}$$

and set of nonzero elements $B_{ts}$ can be defined as:

$$B_{ts} = \{a_{pc} \neq 0 | \; p = 1,2,\ldots.,r_t \; and \; c = 1,2,\ldots,14\}$$

Where, $p = 1,2,\ldots.,r_t$ represents the row and define the pathways and $c = 1,2,\ldots.,14$ represents the column and define the 14 cell-types. A density $D_{ts}$ of nonzero element within matrix $A_{ts}$ can be defined as:

$$D_{ts} = \frac{|B_{ts}|}{14.r_t}$$

The denominator represents total number of elements in the matrix $A_{ts}$. Notably, the density score here relates to how well a theme is represented across a state, rather than the number of different cell-types it can span, which is an important factor in determining a theme's commonality or uniqueness. For this, we included weights $W_{ts}$ for each theme matrix $A_{ts}$ in each state represented by the number of cell-types in which a theme appears

$$W_{ts} = \sum_{q=1}^{14} C_{ts}^{(q)} \ = \begin{cases} 1 \ \ if \ \exists \ p \ such \ that, a_{pq} \neq 0 \\ 0 \ \ if \ \forall \ p, a_{pq} = 0 \end{cases}$$

Here $C_{ts}^{(q)}$ counts $c = 1,2, \ldots.,14$ column of matrix $A_{ts}$ having at least one nonzero element. By including weights, the new density can be calculated as:

$$Z_{ts} = D_{ts}.W_{ts}$$

However, in some instances, $Z_{ts}$ went beyond 1. For it to be $0 \leq Z_{ts} \leq 1$, we calculated the max–min normalization of $Z_{ts}$ in each state $s = 1,2,3,4$ as follows:

$$Z_{ts} = \frac{D_{ts}.W_{ts} - l_s}{L_s - l_s}$$

Where:

$$L_s = max\{D_{ts}.W_{ts} | \ t = 1,2, \ldots.,44\}$$

$$l_s = min\{D_{ts}.W_{ts} | \ t = 1,2, \ldots.,44\}$$

Note that the embedded nature of $A_{ts}$ ensures that the theme density is normalized across all state and themes.

**Disease and Drug enrichment analysis**

Cell-type affected by different diseases-state and drug were determined using hypergeometric overlap analysis (i.e., fisher exact test) implemented by gene-overlap package in R-3.6.0. Given two gene-lists, a significance test for their overlap in comparison with a genomic background representing universe of known genes (21,196 genes, default used by the package) can be described using hypergeometric distribution. The null-hypothesis represents an odd-ratio <1 whereas the alternate hypothesis represents an odd-ratio >1. Disorders and drugs whose gene-sets overlapped significantly with the gene-set of a cell-type were considered related with it. The pool of genes across all diseases and drugs which show significant overlap in a given cell-type represents the vulnerable and druggable transcriptome used for the pathway analysis described above.

**Supplementary Figures (each as separate PDF file) legends and notes:**

**Supplementary Figures 1a & 1b: Characterizing the t-SNE clusters** Each plot depicts the expression (orange) of a known cell type marker (indicated above each plot) across the clusters identified in the single cell RNAseq dataset. Clusters numbers and the complete t-SNE plot are provided in the bottom right for comparison.

**Supplementary Figures 2: Detailed examination of diseases associated with different cell-types.** Heatmap of disease whose gene-sets showed significant (*p-value* < 0.05) overlap with cell-types specific gene sets. Specific diseases are listed at the left. The broader disease categories featured in Figure 3 provided at the right. Cell-type cluster numbers are provided at the top, with PN clusters in purple (left), interneuron clusters in orange (middle), and non-neuronal clusters in blue (right). The color-intensity (light to dark blue) in the heatmap is proportional to -log10(*p-value*), with darker colors indicating more significant overlap between the respective disease and cell-type. Note that these diseases were used to define the disease-state. For a given cell-type the pool of gene driving the significant overlap define the vulnerable transcriptome.

**Supplementary Figures 3: An overview of the approach identifying cell-type specific (control), vulnerable, druggable, and actionable transcriptomes. A)** Upregulated genes from each cell-type cluster were used to identify cell-type specific transcriptomes defining the control state (yellow). **B1)** CNS disease-specific signatures from DisGeNET were used to perform hypergeometric overlap with each cell-type's specific transcriptome. The pool of genes driving the significant overlap (area in blue) for each cell-type cluster were considered vulnerable transcriptomes defining the disease-state. **C)** Similarly, the overlap between cell-type specific transcriptomes and signatures of drugs with known mechanism of action (MOA) were used to identify druggable transcriptomes (area in maroon) defining the druggable state. **B2)** To identify actionable transcriptome defining the drug's action in the disease-state or the actionable state, the signatures of drugs with known MOA were overlapped with vulnerable transcriptome (area in green). Transcriptomes for each state were used to perform pathway analysis using the gene ontology database. The significant pathways ($q$-*value* < *0.05*) for each state were organized into biological themes which were quantitatively compared between different states using density scores. The rectangle defines the universe (U) of genes used as background to perform hypergeometric analysis. See the methods section for further details.

**Supplementary Figures 4: Drugs and MOAs linked to druggable and actionable states in various cell types. A)** Percent of Drugs (with known MOA) associated with druggable (maroon) and actionable-state (green) in different cell-type clusters. Cluster numbers are provided at the left, with PN clusters in purple, interneuron clusters in orange, and non-neuronal clusters in blue. **B)** Percentage of specific mechanisms of action (MOA) associated with PNs, interneurons (INT), and non-neurons (NN) in druggable- (maroon) and actionable—state (green). MOAs (left labels) as defined by the connectivity map and comparative toxicogenomics database (see methods) are used. The lighter-to-darker shades of blue represent the percentage of MOAs associated with each cell class. The number of drugs for each MOA is provided in parentheses. See Supplementary Table 9 for more details.

**Supplementary Figures 5: Cell-type specific functional comparison between control and druggable state.** Heatmap of significant ($q$-*value* <0.05) pathways for each cell-type specific gene-set and druggable transcriptome representing control (yellow) and druggable state (red), respectively. Cell-type cluster numbers are provided at the top, with PN clusters in purple (left), interneuron clusters in orange (middle), and non-neuronal clusters in blue (right). Pathways are organized by functional themes provided at the left. Each state's theme-densities are represented by solid circles of their respective colors in the heatmap. An increase in druggable-state density (marked with red vectors) represent themes involving additional cell-type or pathways during druggable state. See Supplementary Table 12 & 13 for more details.

**Supplementary Figures 6: Cell-type specific functional comparison between control and actionable state.** Heatmap of significant (*q-value* <0.05) pathways for each cell-type specific gene-set and actionable transcriptome representing control (yellow) and actionable (drug action in disease sate) state (green), respectively. Cell-type cluster numbers are provided at the top, with PN clusters in purple (left), interneuron clusters in orange (middle), and non-neuronal clusters in blue (right). Pathways are organized by functional themes provided at the left. Each state's theme-densities are represented by solid circles of their respective colors in the heatmap. An increase in actionable-state density (marked with red vectors) represent themes involving additional cell-type or pathways during actionable -state. See Supplementary Table 12 &13 for more details.

**Supplementary Figures 7: Cell-type specific functional comparison between druggable and actionable state.** Heatmap of significant (*q-value* <0.05) pathways for druggable and actionable transcriptome representing druggable (red) and actionable state (green), respectively. Cell-type cluster numbers are provided at the top, with PN clusters in purple (left), interneuron clusters in orange (middle), and non-neuronal clusters in blue (right). Pathways are organized by functional themes provided at the left. Each state's theme-densities are represented by solid circles of their respective colors in the heatmap. An increase in actionable-state density (marked with red vectors) represent drug effects that emerge in the presence of disease. **Note** that the increased involvement of deep layer PNs (cluster 13) with actionable state (Supplementary Figure 4) is associated with *glutamate*, *GPCR signaling*, synaptic changes (*modulation of chemical synaptic transmission, presynapse and transsynaptic signaling*), axon, dendrites and cell surface receptor signaling pathway. Whereas those involving across layer PNs (cluster 00) were involved in *homeostatic process*, *calcium,* and *regulation of protein phosphorylation.* See Supplementary Table 12 &13 for more details.

**Supplementary Figures 8: Enrichment of multiple cell-type in a GO term represent their joint participation in the functional pathway represented by the that GO term. Top:** A GO term represents set of coordinated genes (green lines) linked to a biological-process, molecular-function, or cellular-component. Their overlap with various microcircuitry cell-types (colored circles), as defined by unique gene-sets, varies with different biological conditions, for instance aging. **Bottom:** Cell types sharing the GO-terms share a common input. For instance, *Vip* and pyramidal neurons share GO-terms related to post-synaptic processes, thus highlighting their common afferent input which alters with age[13, 14]. Colored circles in the top correspond to colored neuronal population below. ***VIP*****:** vasoactive intestinal peptide expressing interneurons; ***SST*****:** Somatostatin expressing interneurons; ***PV*****:** Parvalbumin expressing interneurons and ***PYC*****:** Pyramidal neurons. Figure adapted from[15].

**Supplementary Tables (each as separate excel worksheet), legends and notes:**

**Supplementary Table 1 [sheet: ST1]: Seurat output and filtering of 2-fold markers.** Output of sn-RNAseq analysis performed using Seurat package in R. Each gene is associated with a cell-type cluster (named 0 through 13). The output has following column headers:
**P_val:** unadjusted p-value of a genes enrichment in a given cluster compared to the remaining clusters
**p_val_adj:** adjusted p-value, based on Bonferroni correction using all features in the dataset.
**avg_logFC:** and average log fold change in expression associated with its native cluster (pct.1) compared to other clusters (pct.2).
**pct.1:** Percentage of cells where the genes are detected in the native cluster
**pct.2:** Percentage of cells where the genes are detected in the remaining other clusters
**Frequency:** Number of clusters in which the gene is significantly enriched
**MarkerSelection:** genes which are minimum two times more enriched in pct.1 than pct.2
**Ratio:** pct.1/pct.2

**Supplementary Table 2 [sheet: ST2]: List of 2-fold enriched and upregulated genes in each cell type cluster.** The list represents the control state and was used to perform the GO analysis.

**Supplementary Table 3 [sheet: ST3]: GO analysis output for each cell-type specific gene-list (from ST2) representing the control state.** The output has following column headers:
**State:** For control state
**GO Category:** different gene-ontology category to which the pathways belong. CC: Cellular Component, BP: Biological Process and MF: Molecular Function.
**Cell-type Cluster ID:** Cell-type cluster to which the input gene-list belongs
**GO Term:** Gene ontology pathways with their respective IDs.
***Homo sapiens* – REFLIST (20851):** Number of genes from the Human reference gene (i.e., out of 20851) list that map to a given pathway.
**Upload_1:** Number of genes in the predictive gene list (up: 62 and down: 59) that map to a given pathway.
**Expected:** Based on the *Homo sapiens* – REFLIST, the number of genes expected in the cell-type specific gene list for a given pathway.
**Fold Enrichment:** Number in column "Upload_1 (up: 62)/(down:59)" divided number in column "Expected", i.e., fold enrichment of genes observed in predictive gene list (up and down separately) over the expected value.
**P-value:** Raw p-value as determined by Fisher's exact test.
**FDR:** Benjamini-Hochberg adjusted p-value.
**ES:** Enrichment score = -log10(FDR corrected P-value)

**Supplementary Table 4 [sheet: ST4]:** We examined the enrichment of disease specific gene sets obtained from DisGeNET (https://www.disgenet.org/) in the cell-specific gene sets (ST2). Gene-sets form 112 disorders (rows) were found enriched across the all the ACC cell-types (column B through O). The value (in blue) represents the enrichment score [ES = $-\log_{10}$(*p*-

*value*)] of each disease in their respective cell-type. Abridged disease categories as provided by DisGeNET is shown in column P.

**Supplementary Table 5 [sheet: ST5]:** List of vulnerable transcriptomes in each cell type cluster. The list represents the disease state and was used to perform the GO analysis.

**Supplementary Table 6 [sheet: ST6]:** GO analysis output for each cell-type specific vulnerable transcriptome (from ST5) representing the disease state. The output column header details same as ST3.

**Supplementary Table 7 [sheet: ST7]:** List of druggable transcriptomes in each cell type cluster. The list represents the druggable state and was used to perform the GO analysis.

**Supplementary Table 8 [sheet: ST8]:** List of actionable transcriptomes in each cell type cluster. The list represents the druggable state and was used to perform the GO analysis.

**Supplementary Table 9 [sheet: ST9]:** We examined the enrichment of drug [with known mode of action or target, (column 'AL' and 'AM', respectively)] specific gene sets obtained from DSigDB (http://dsigdb.tanlab.org/DSigDBv1.0/) in the cell-type specific vulnerable- and druggable- transcriptome (from ST5 and ST7, respectively). The values (in green) represent the enrichment score [ES = $-\log_{10}(q\text{-}value)$] of each drug in their respective cell-type for a given state.

**Supplementary Table 10 [sheet: ST10]:** GO analysis output for each cell-type specific druggable transcriptome (from ST7) representing the druggable state. The output column header details same as ST3.

**Supplementary Table 11 [sheet: ST11]:** GO analysis output for each cell-type specific actionable transcriptome (from ST8) representing the actionable state. The output column header details same as ST3.

**Supplementary Table 12 [sheet: ST12]:** Comparison of complete GO profile (rows) across all cell-types and state (column) created first by vertically stacking the pathway profiles provided in ST3, ST6, ST10 and ST11 and then pivoting them. The value represents the enrichment score of each pathway in their respective cell-type in a given state.

**Supplementary Table 13 [sheet: ST13]:** Filtering of GO profiles from ST12 based on themes provided in column "BK".
