## Supplementary figures and images for "Differential Vulnerability of Anterior Cingulate Cortex Cell-Types to Diseases and Drugs"

### Supplementary Figure 1a

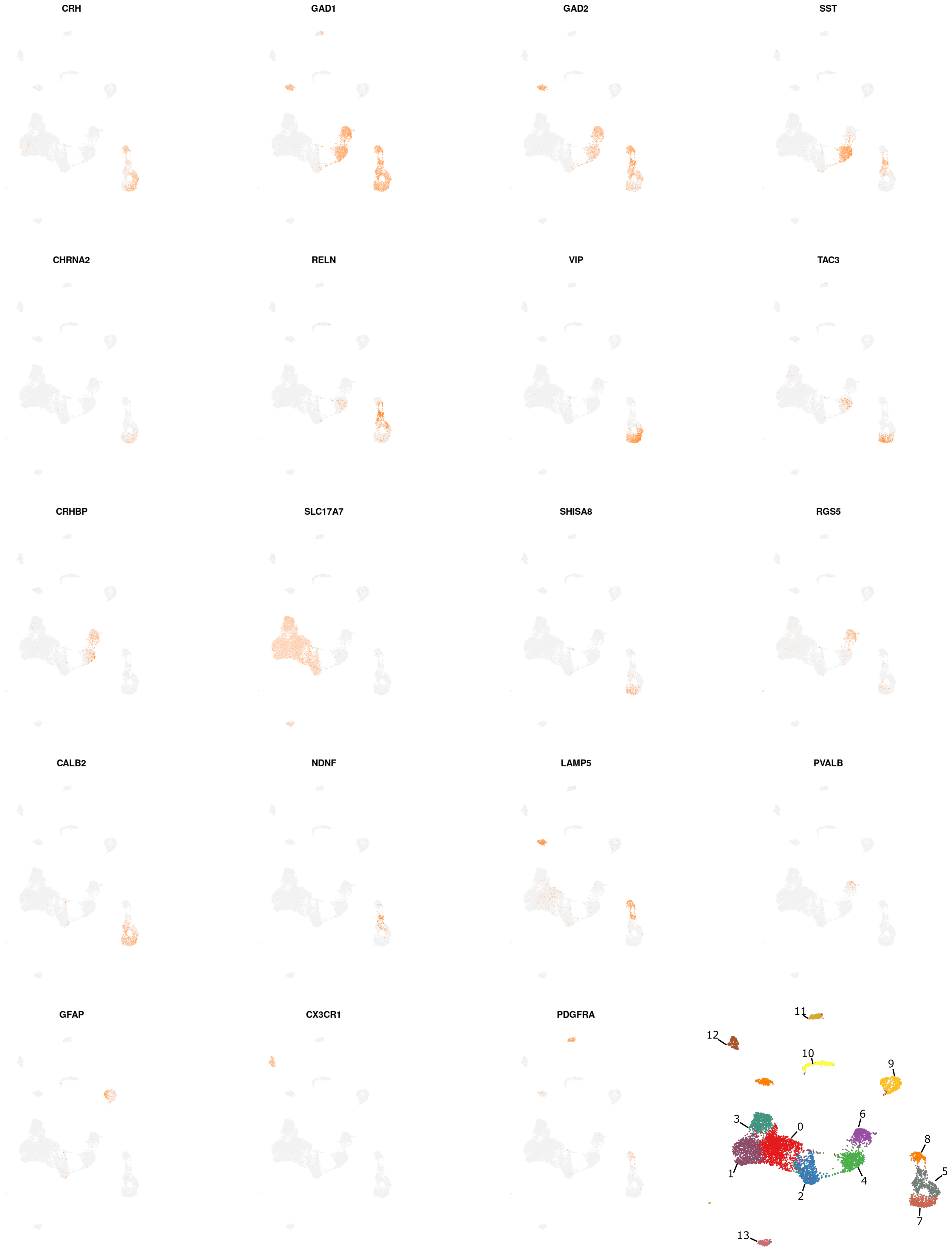


Supplementary Figure 1a

### Supplementary Figure 1b

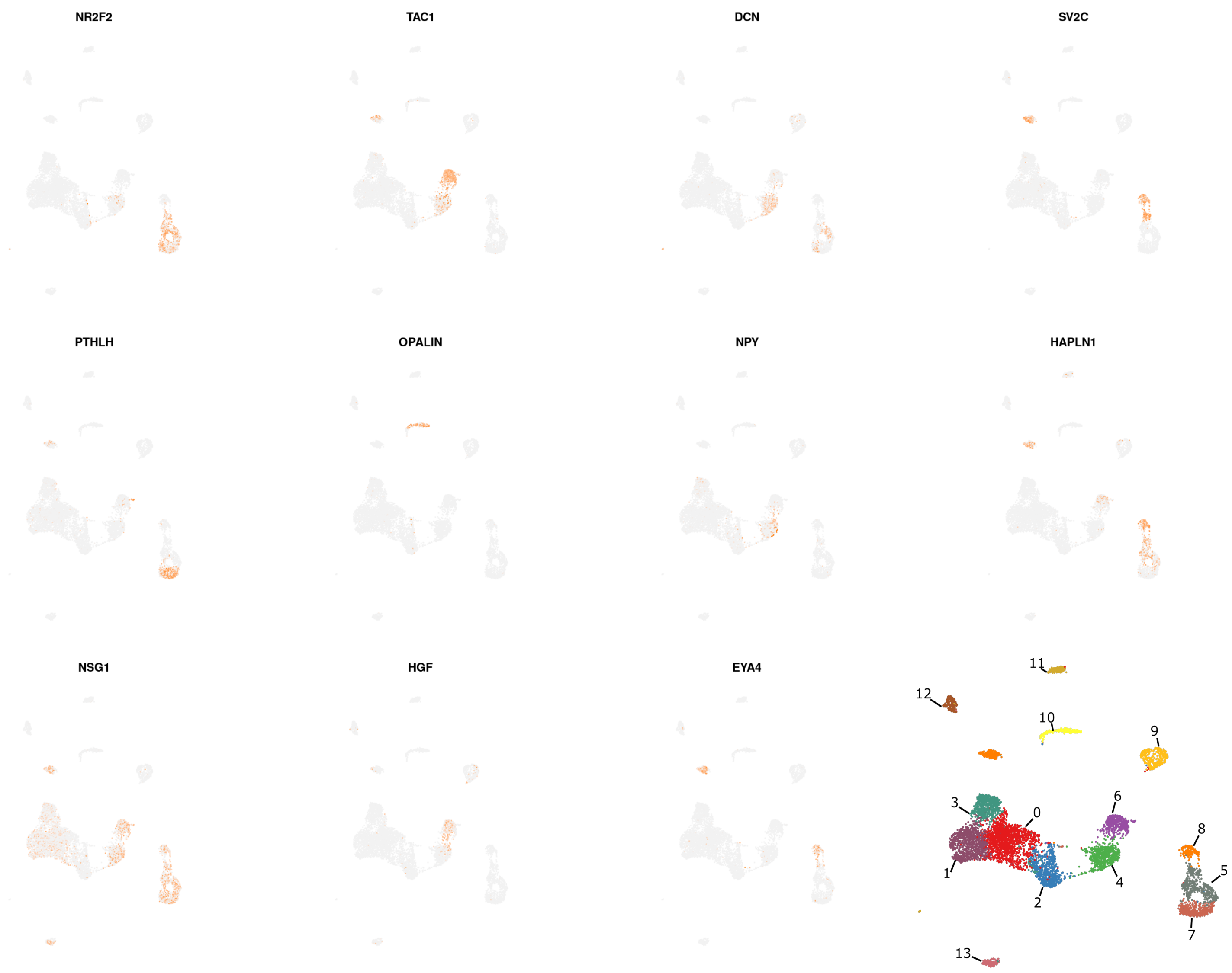


#### Supplementary Figure 1b

### Supplementary Figure 2

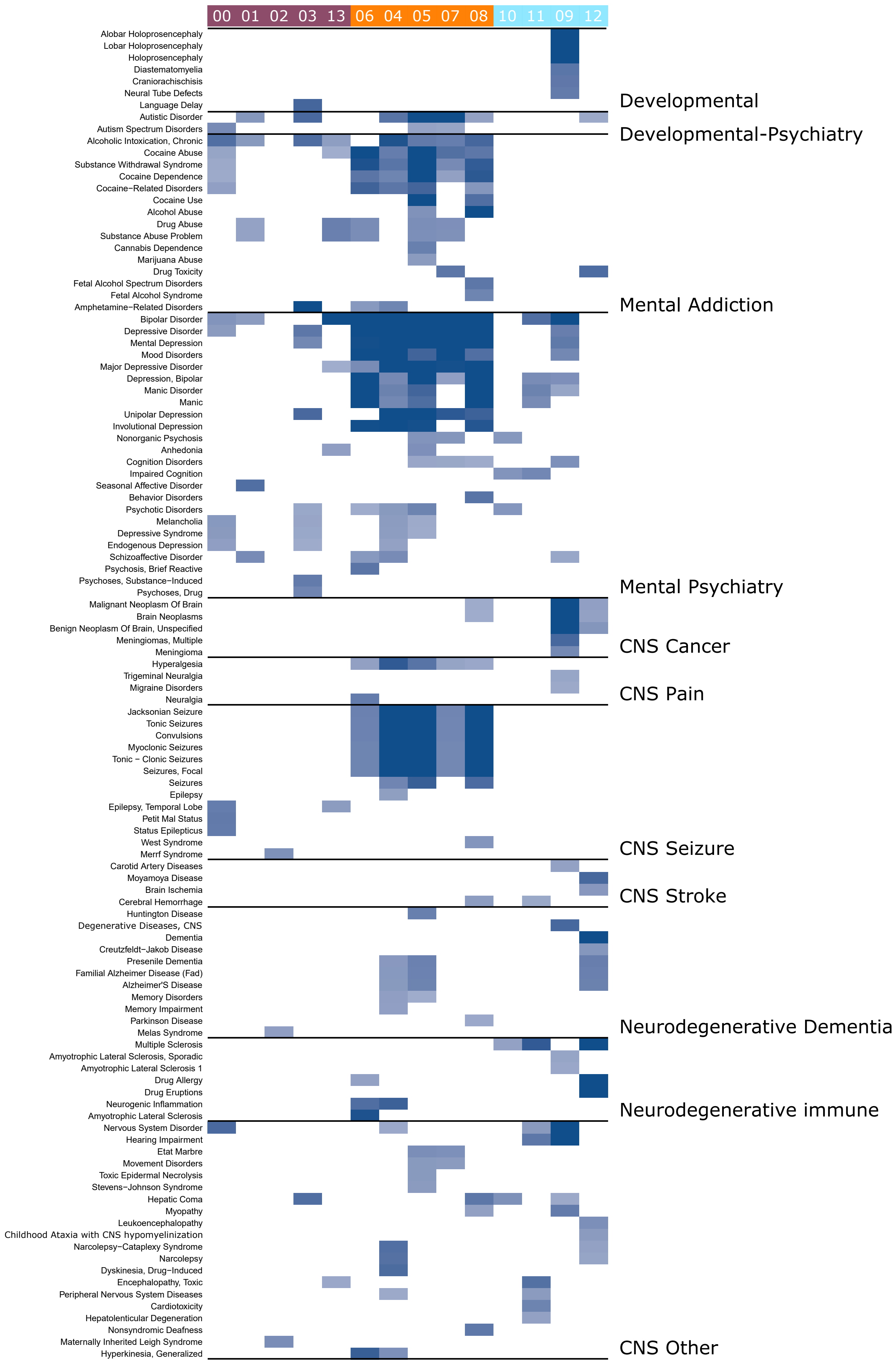


#### Supplementary Figure 2

### Supplementary Figure 3

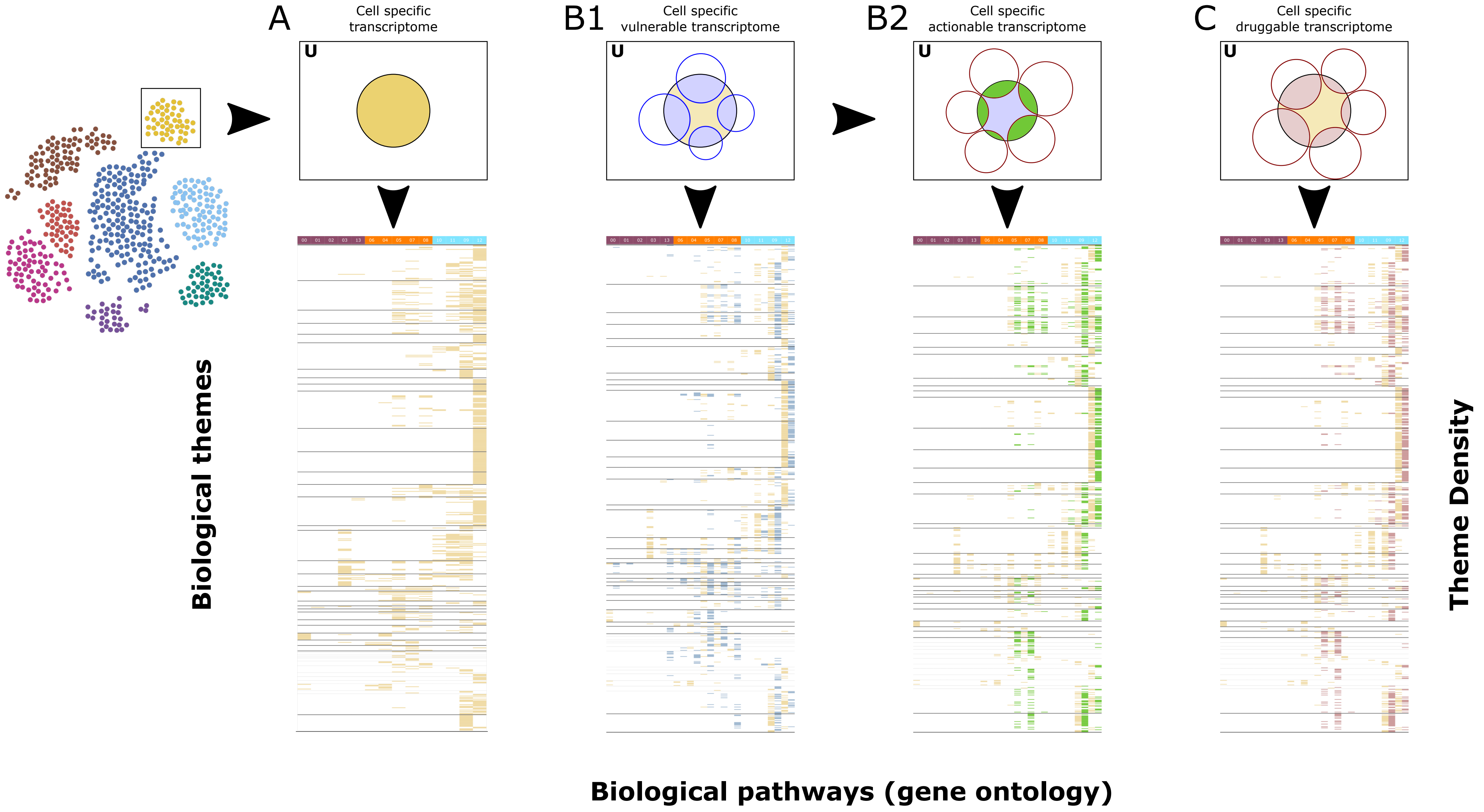


Supplementary Figure 3

### Supplementary Figure 5

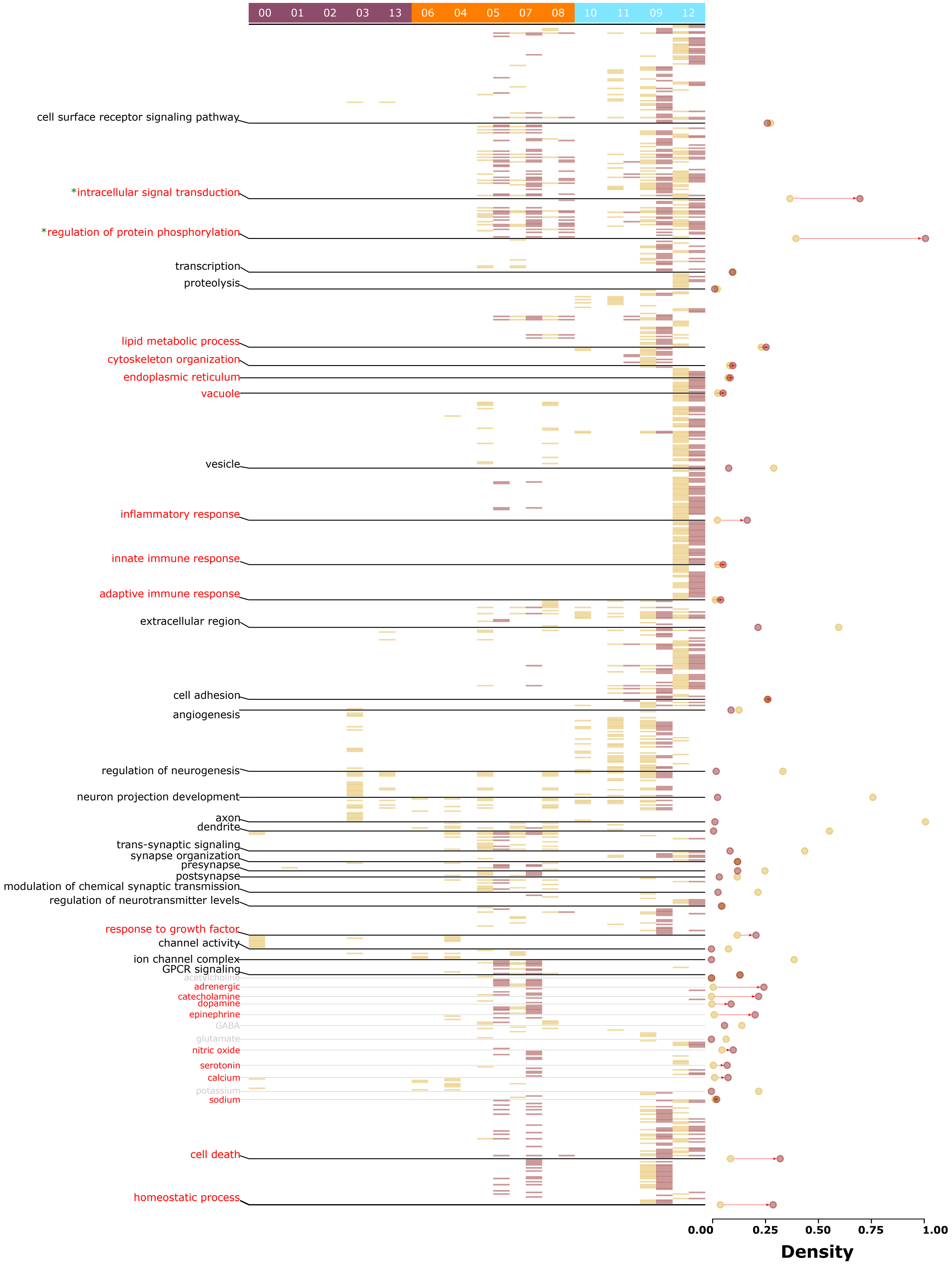


#### Supplementary Figure 5

### Supplementary Figure 6

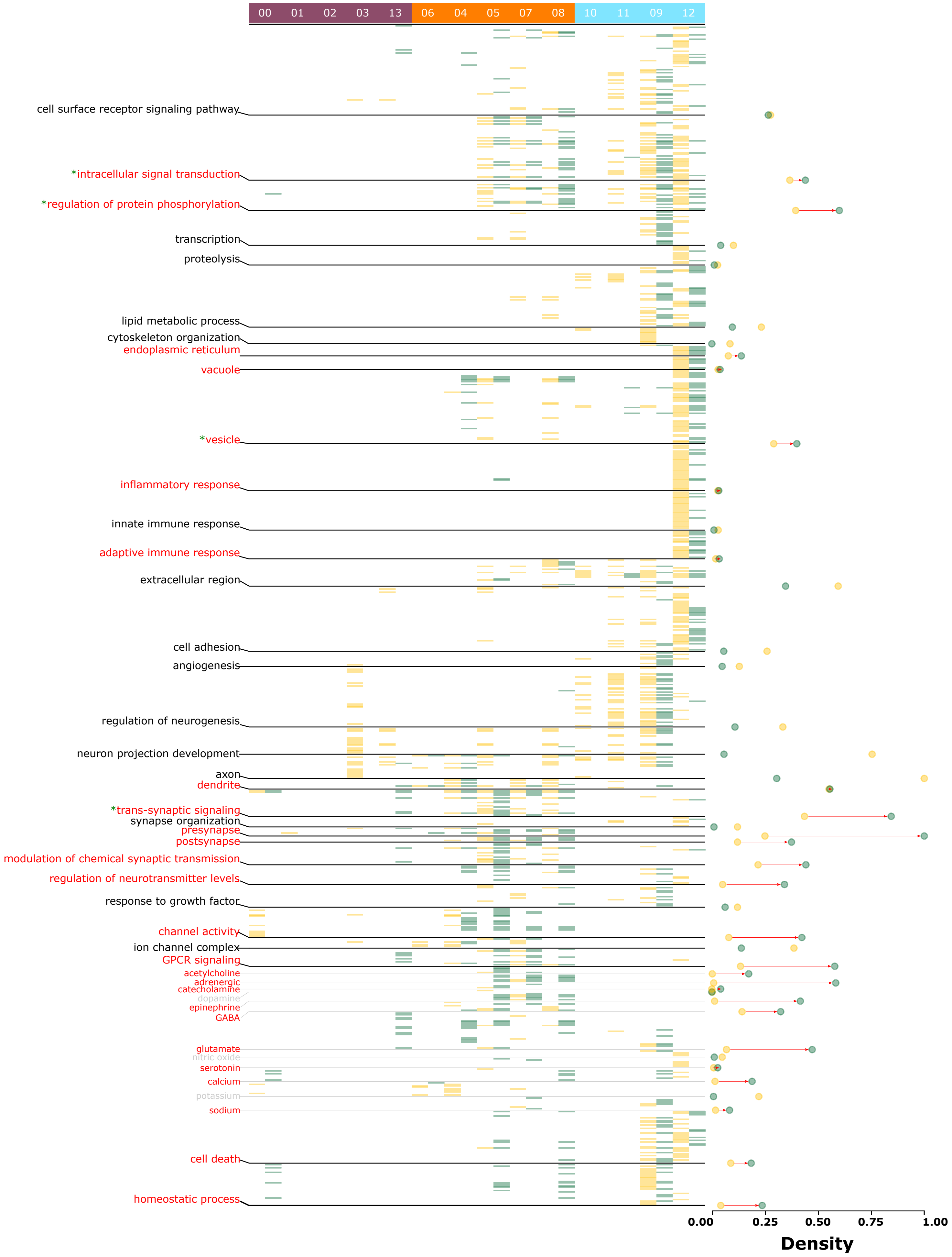


#### Supplementary Figure 6

### Supplementary Figure 7

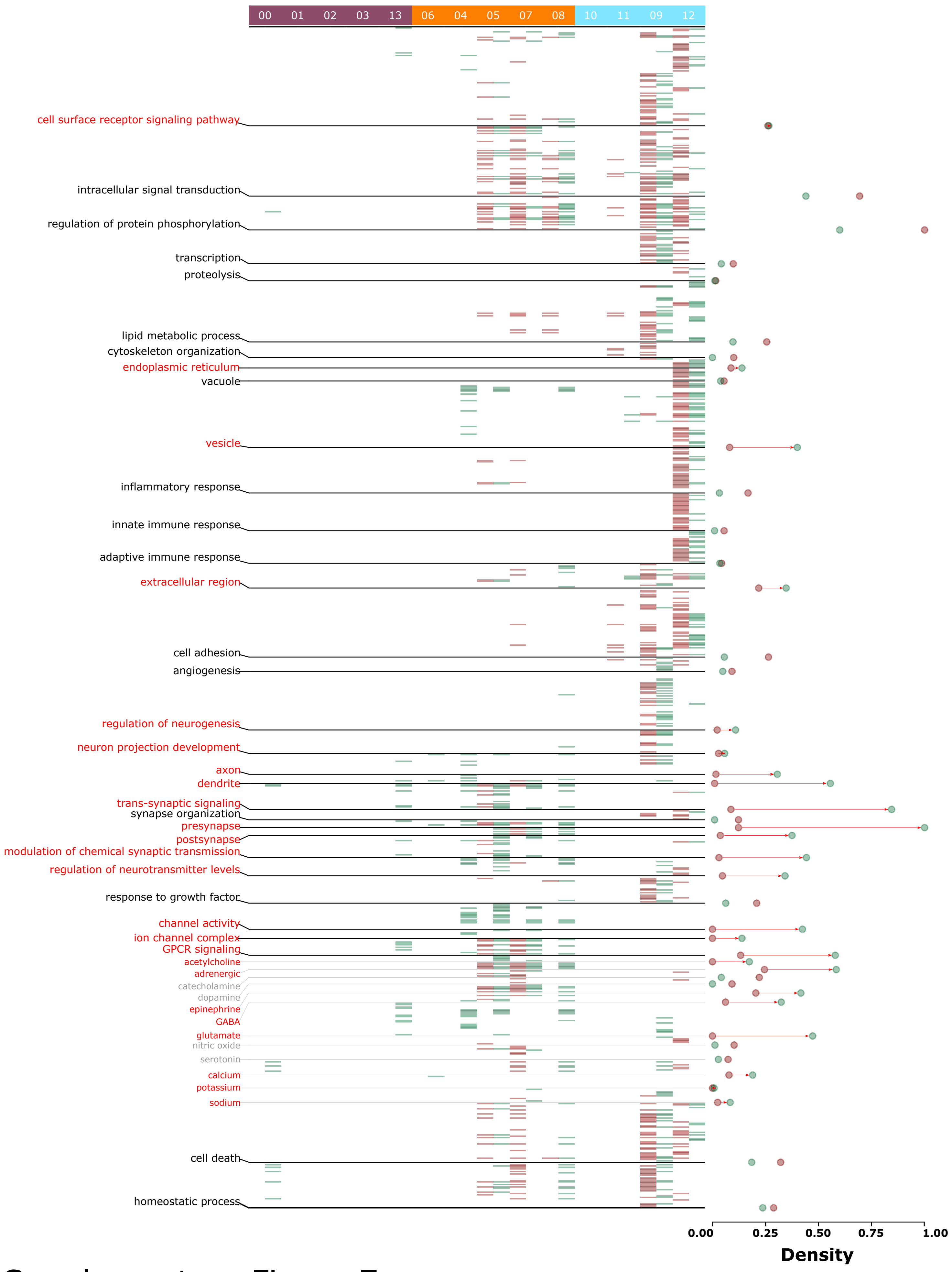


Supplementary Figure 7

### Supplementary Figure 8

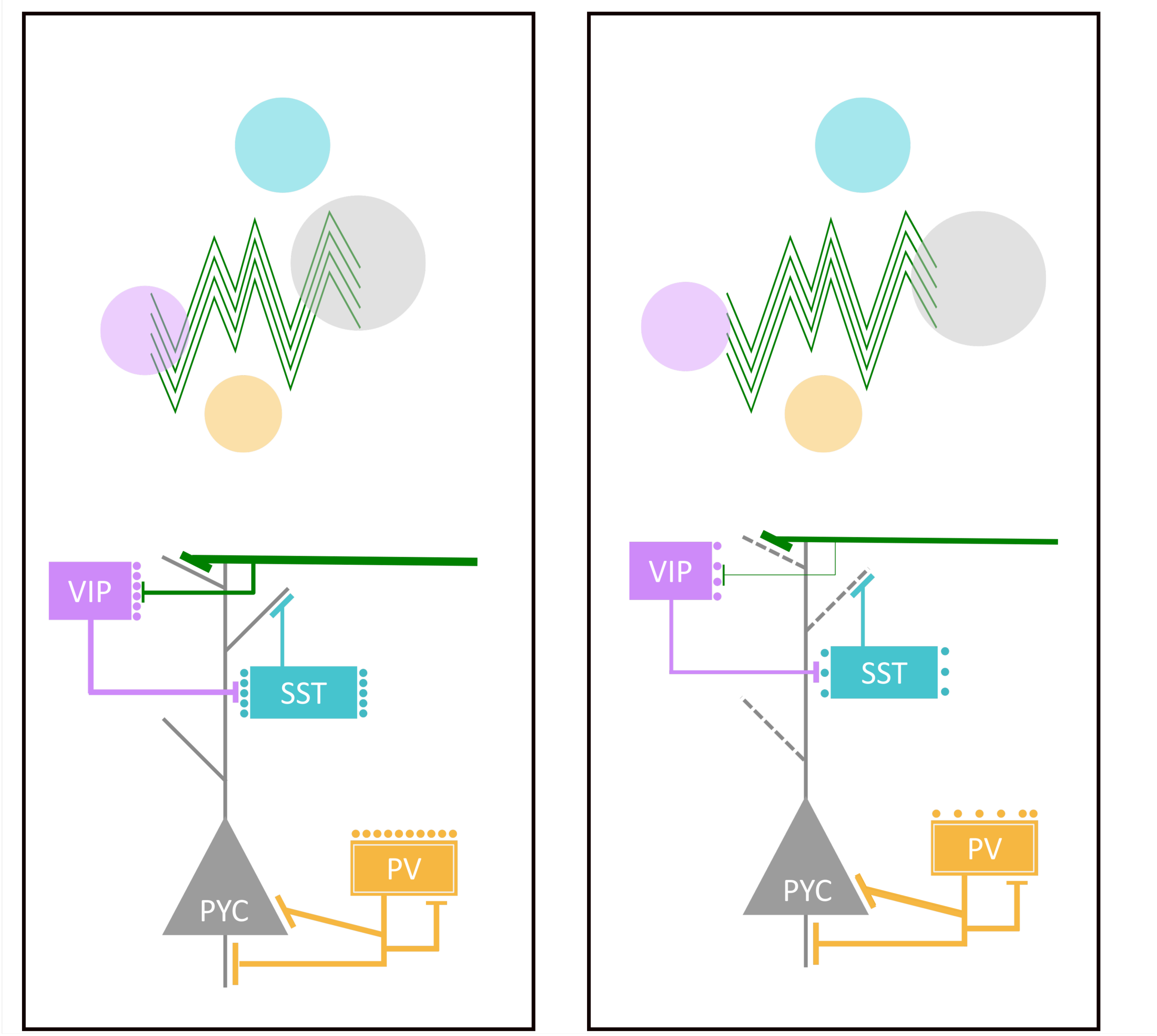


Supplementary Figure 8
