## Supplementary Figure 4 for "Differential Vulnerability of Anterior Cingulate Cortex Cell-Types to Diseases and Drugs"

A

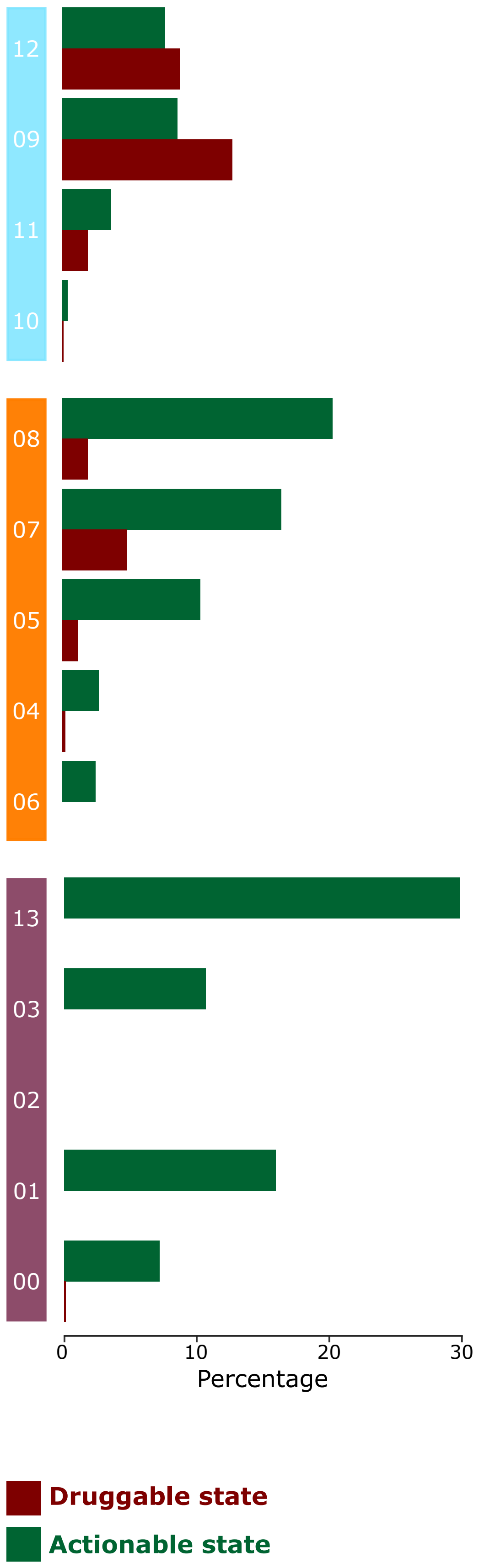


B

| | PN | INT | NN | PN | INT | NN |
|---|---|---|---|---|---|---|
| ABL inhibitor (5) | | 80.0 | | | | 20.0 |
| Acetylcholine receptor agonist (6) | 50.0 | 33.3 | 16.7 | | | |
| Acetylcholine receptor antagonist (8) | 75.0 | 37.5 | 25.0 | | 12.5 | 25.0 |
| Adenosine receptor antagonist (7) | 42.9 | 14.3 | 42.9 | | | 42.9 |
| Adrenergic receptor agonist (28) | 39.3 | 50.0 | 32.1 | | 17.9 | 28.6 |
| Adrenergic receptor antagonist (40) | 45.0 | 55.0 | 25.0 | | 20.0 | 22.5 |
| Androgen receptor antagonist (7) | 85.7 | 14.3 | | | | |
| Angiotensin receptor antagonist (8) | 12.5 | | 75.0 | | | 50.0 |
| ATPase inhibitor (18) | 33.3 | 22.2 | 55.6 | | | 16.7 |
| BCR-ABL kinase inhibitor (10) | 40.0 | 90.0 | 30.0 | | 30.0 | 30.0 |
| Benzodiazepine receptor agonist (8) | 50.0 | 37.5 | 37.5 | | 12.5 | 12.5 |
| Calcium channel blocker (21) | 57.1 | 33.3 | 23.8 | | 9.5 | 19.0 |
| Carbonic anhydrase inhibitor (5) | 80.0 | 80.0 | | | 20.0 | |
| Cyclooxygenase inhibitor (23) | 43.5 | 13.0 | 69.6 | | | |
| Cytochrome P450 inhibitor (9) | 55.6 | 11.1 | 33.3 | | 11.1 | 22.2 |
| Dihydrofolate reductase inhibitor (5) | 80.0 | | 20.0 | | | |
| DNA replication inhibitor (5) | 60.0 | 20.0 | | | | 20.0 |
| DNA synthesis inhibitor (5) | 80.0 | 20.0 | | | | |
| Dopamine receptor agonist (15) | 73.3 | 40.0 | 40.0 | | 6.7 | 20.0 |
| Dopamine receptor antagonist (67) | 82.1 | 40.3 | 7.5 | | 6.0 | 9.0 |
| EGFR inhibitor (18) | 27.8 | 88.9 | 33.3 | | 5.6 | |
| Estrogen receptor agonist (14) | 71.4 | 21.4 | 21.4 | | | |
| Estrogen receptor antagonist (12) | 50.0 | 41.7 | 25.0 | | | 33.3 |
| FLT3 inhibitor (8) | 37.5 | 75.0 | 50.0 | | 12.5 | 12.5 |
| Glucocorticoid receptor agonist (43) | 30.2 | 32.6 | 30.2 | | 9.3 | 39.5 |
| HDAC inhibitor (6) | 33.3 | | 50.0 | | | 50.0 |
| Histamine receptor antagonist (16) | 50.0 | 31.3 | 18.8 | | | 25.0 |
| HMGCR inhibitor (11) | 18.2 | 9.1 | 63.6 | | 9.1 | 36.4 |
| Insulin sensitizer (10) | 40.0 | 40.0 | 40.0 | | | 30.0 |
| MTOR inhibitor (5) | 40.0 | 20.0 | 20.0 | | 20.0 | 40.0 |
| Phosphodiesterase inhibitor (17) | 76.5 | 23.5 | 35.3 | | 5.9 | 11.8 |
| PKC inhibitor (6) | 50.0 | 50.0 | 16.7 | | | 16.7 |
| Protein synthesis inhibitor (9) | 22.2 | 33.3 | 22.2 | | 11.1 | 44.4 |
| RAF inhibitor (5) | | 60.0 | 60.0 | | 40.0 | |
| Retinoid receptor agonist (6) | 33.3 | 16.7 | 66.7 | | | 66.7 |
| Selective serotonin reuptake inhibitor (SSRI) (6) | 83.3 | 16.7 | | | | |
| Serotonin receptor agonist (17) | 76.5 | 52.9 | 11.8 | | 23.5 | 5.9 |
| Serotonin receptor antagonist (12) | 83.3 | 75.0 | | | 16.7 | |
| Topoisomerase inhibitor (11) | 18.2 | 27.3 | 36.4 | | | 45.5 |
| Tubulin inhibitor (10) | 30.0 | 20.0 | 40.0 | | | 40.0 |

Supplementary Figure 4
